# Comparative phylogenomic patterns in the Baja California avifauna, their conservation implications, and the stages in lineage divergence

**DOI:** 10.1101/2021.08.18.456789

**Authors:** Hernán Vázquez-Miranda, Robert M. Zink, Brendan J. Pinto

**Affiliations:** Colección Nacional de Aves, Departamento de Zoología, Instituto de Biología, Universidad Nacional Autónoma de México, Ciudad de México, C.P. 04510, Mexico; School of Natural Resources, School of Biological Sciences and Nebraska State Museum, University of Nebraska, Lincoln, NE 68583, USA; Center for Evolutionary Medicine & Public Health, Arizona State University, Tempe, AZ 85281, USA; Department of Zoology, Milwaukee Public Museum, Milwaukee, WI 53233, USA

**Keywords:** Lineage divergence, phylogeography, genotype-by-sequencing, RADseq, population structure, conservation, Baja California

## Abstract

Comparative phylogeography explores the historical congruence of co-distributed species to understand the factors that led to their current genetic and phenotypic structures. Even species that span the same biogeographic barrier can exhibit different phylogeographic structures owing to differences in effective population sizes, marker bias, and dispersal abilities. The Baja California peninsula and adjacent desert regions include several biogeographic barriers that have left phylogeographic patterns in some but not all species. We found that mitochondrial DNA, single nuclear genes, and genome-wide SNP data sets show largely concordant phylogeographic patterns for four bird species along the Baja California peninsula and adjacent mainland: cactus wren, Gila woodpecker, California gnatcatcher, and LeConte’s thrasher. The cactus wren and LeConte’s thrasher show a concordant historical division at or near the Vizcaíno Desert in north-central Baja California, the Gila woodpecker appears to be at an intermediate stage of divergence, and the California gnatcatcher lacks notable phylogeographic structure. None of these four species are classified taxonomically in a way that captures their evolutionary history with the exception of the LeConte’s thrasher. We also present mtDNA data on small samples of ten other species that span the Vizcaíno Desert, with five showing no apparent division, and five species from the Sierra de la Laguna, all of which appear differentiated. Reasons for contrasting phylogeographic patterns should be explored further with genomic data to test the extent of concordant phylogeographic patterns. The evolutionary division at the Vizcaíno desert is well known in other vertebrate species, and our study further corroborates the extent, profound effect and importance of this biogeographic boundary. The areas north and south of the Vizcaíno Desert, which contains considerable diversity, should be recognized as historically significant areas for conservation.

## 1. Introduction

In theory, species existing over the same area ought to have experienced the same geological history and display concordant phylogeographic patterns (Bermingham and Mortiz, 1998). However, if some species are recent arrivals to the region, differ in their ability to disperse across biogeographic barriers, or possess different effective population sizes then phylogeographic patterns may differ (Zink, 2002, 2010; Harrington et al., 2018). As a result, co-distributed species can exhibit shallow, strong, or no divergence across a common geographic area. Comparison of multiple co-distributed species, or comparative phylogeography, lends insight into factors that shape diversity (Bermingham and Moritz, 1998; Zink, 1996; Avise et al., 2016) and can help identify geological or ecological factors causing divergence, or the lack thereof. Furthermore, these comparisons can inform our understanding of speciation, because depending on the species concept, lineage divergence can be considered one and the same process (Roux et al., 2016). In addition, the use of different molecular markers can help elucidate whether a concordant or discordant pattern can also be due to factors such as sex-biased dispersal, marker ploidy, or genetic incompatibilities (Hill et al., 2015). Baja California and the associated arid lands of North America have provided important areas for exploring comparative phylogeographic patterns (Scheinvar et al., 2020; Gamez et al., 2017). Many studies (e.g., Dolby et al., 2015) have implicated the existence of a mid-peninsular seaway at about the latitude of 27°–30°N (Riddle et al., 2000; Lindell et al., 2006), although the current patterns of annual temperature and rainfall do not suggest a discrete ecological difference at present (Fig. S1).

The avifauna of the Baja California peninsula includes species with contrasting patterns of genetic differentiation at a minimum of two biogeographic scales (Erickson et al., 2013), which present several testable hypotheses. For example, several differentiated avian taxa occupy the Sierra de la Laguna, isolated mountains at the southern end of the Baja peninsula (Garrick et al., 2009; Morrone, 2021), including the San Lucas robin (*Turdus migratorius confinis* or *T*. *confinis*), acorn woodpecker (*Melanerpes formicivorus angustifrons*; Honey-Escandón et al., 2008), white-breasted nuthatch (*Sitta carolinensis lagunae*; Spellman and Klicka, 2007; Walstrom et al., 2012), bushtit (*Psaltriparus minimus*), and Baird’s junco (*Junco bairdi*; Friis et al., 2016). The divergence of these taxa is possibly a result of an isthmus (“La Paz”) that separated the cape district from the rest of the peninsula (Riddle et al., 2000; Dolby et al., 2015). Several studies have suggested that these cape district endemics have their nearest relatives on the Mexican mainland (Friis et al., 2016; Spellman and Klicka, 2007; Honey-Escandón et al., 2008).

Another well-studied biogeographic gap is located at the approximate latitude of the Vizcaíno desert (Dolby et al. 2015; Fig. 1; Table S1). Multiple avian taxa, including the LeConte’s thrasher (*Toxostoma lecontei*), cactus wren (*Campylorhynchus brunneicapillus*), verdin (*Auriparus flaviceps*) and Gila woodpecker (*Melanerpes uropygialis*) appear to exhibit phylogenetic divisions at or near the Vizcaíno Desert (Zink et al., 1997, 2001; Vázquez-Miranda, 2014; Vázquez-Miranda et al., 2017), a pattern found in many other species (Riddle et al., 2000a; Peterson et al., 2013). However, the phylogeographic breaks are not all consistent. For example, Harrington et al. (2018) found a different genetic division in a rattlesnake, *Crotalus ruber*, centered south of the Vizcaíno Desert and they concluded there was secondary contact with northward gene flow. These patterns also contrast with other species, such as the California gnatcatcher (*Polioptila californica*), for which prior data show no apparent divisions across Baja (Zink et al., 2000, 2013). Thus, there are at least two testable phylogeographic structures found in Baja California relating to the mid-peninsular area near the Vizcaíno Desert and the potential seaway that might have affected co-distributed species.

**Figure 1:**
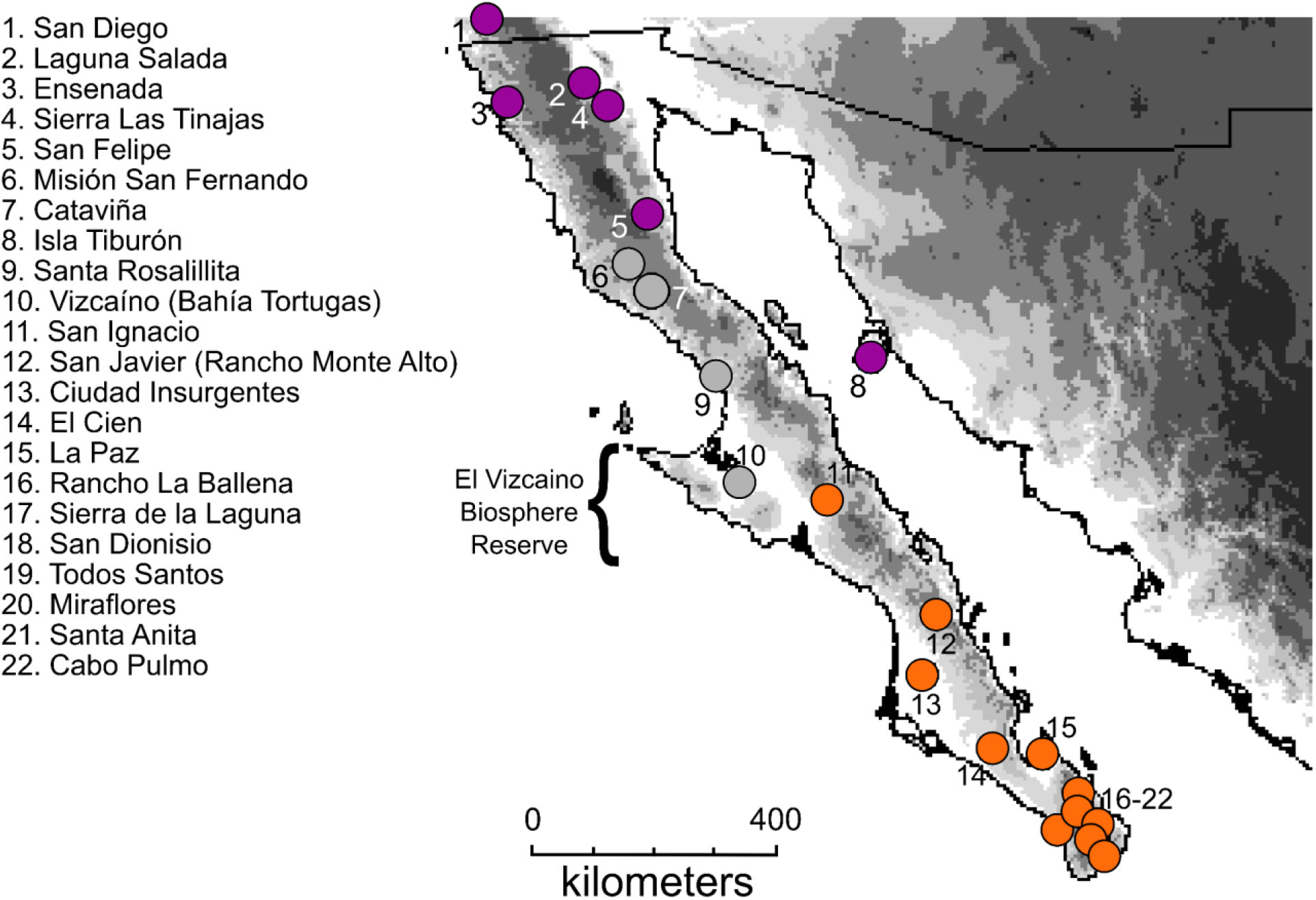
Location of samples north (darker/purple) or south (lighter/orange) of the Vizcaíno Desert. Samples from elsewhere to the north and east not shown. Sample sites that appear to differ between species are shown as intermediate/gray.

Choice of molecular markers might influence the power of tests of comparative phylogeographic hypotheses (Ballard and Whitlock, 2004; Edwards et al., 2000; Leavitt et al., 2017). In the LeConte’s thrasher, Vázquez-Miranda et al. (2017) showed that 16 nuclear genes (nuDNA) recovered the same history as that inferred from mitochondrial genes (mtDNA). Similarly, in the California Gnatcatcher a dataset consisting of eight nuDNA loci corroborated earlier phylogeographic findings using only mtDNA (Zink et al., 2000, 2013, 2016). However, it is possible that few molecular markers or sampling biases (from using genes that have been historically used in phylogenetic studies) might inhibit detection of relatively shallow evolutionary divisions. Importantly, since the advent of population genomic technologies there have been cases in which population-level genomic (single nucleotide polymorphism; SNP) data have been used, both to confirm and reject work based on only a few loci (e.g., Pedraza-Marrón et al., 2019; Pinto et al., 2019; Ramírez-Reyes et al., 2020). Thus, an open question is whether genome-scale analyses will reproduce the same patterns identified previously in these avian taxa from the desert southwest and Baja California.

Here, we report the results of genomic SNP data [genotype-by-sequencing (GBS)] of geographic variation in cactus wren, Gila woodpecker, California gnatcatcher, and LeConte’s thrasher. In addition, due to its potential conservation implications, we generated an additional dataset of restriction site-associated sequencing (RADseq) aggregating independent loci for the California gnatcatcher. We present these new SNP data alongside previously unpublished analyses of mitochondrial and nuclear genes for the cactus wren and Gila woodpecker. Additionally, we conduct preliminary analyses using mtDNA data for an additional 10 species taken on either side of the Vizcaino desert and five species sampled from the Sierra de la Laguna. Our main goal in this study is to identify if there are corresponding patterns in mtDNA, nuDNA, and genome-wide data for these four main study species and exploring the biogeographic gaps at the latitude of the Vizcaíno desert (Riddle et al., 2008; Peterson et al., 2013).

### 1.1 Review of taxa and prior nuDNA loci and mtDNA studies

- Cactus wren: Zink et al. (2001) reported a phylogeographic division at approximately 30°N, based on a small amount of sequence (298 bp) from the mtDNA control region. Vázquez-Miranda (2014) sequenced the mitochondrial ND2 gene (1041 bp, 9 individuals north of and 10 south of the Vizcaíno Desert) and six nuDNA loci (1 Z-linked, 5 autosomal; 3191 bp total, average samples sizes 10 individuals each north and south of the Vizcaíno). Analyses of these nuDNA data recovered the same phylogeographic structure (Fig. S2) that was identified previously (Zink et al., 2001). The timing of divergence was estimated at 2.3 million years before present (Vázquez-Miranda, 2014). In a phylogenetic analysis of *Campylorhynchus* that incorporated two mtDNA genes and 22 nuDNA loci (12 Z-linked and 10 autosomal), Vázquez-Miranda and Barker (2021) found the same north-south split. An ecological niche model for the LGM distribution suggested two allopatric groups (Zink, 2014) north and south of the Vizcaíno desert. Thus, for the cactus wren, previous results from the nuDNA loci and mtDNA were concordant and showed a distinct phylogeographic split at the implied location of the ancient mid-peninsular seaway (latitude 27°–30°N).
- Gila woodpecker: Vázquez-Miranda (2014) sequenced the ND2 gene (1041 bp) for 41 individuals north and 20 south of the Vizcaíno Desert, one Z-linked locus (1044 bp; 54 individuals north, 32 south) and three autosomal loci (1866 bp; 54 individuals north and 32 south for Fib7, 6 and 8 north and 10 south for GAPDH and TGFB2, respectively). The mtDNA gene tree (Fig. S3) of 61 individuals is nearly reciprocally monophyletic for samples on either side of the Vizcaíno desert (an individual (BCHVM097) from Cataviña occurs in the southern group whereas four occur in the northern group); the two groups are separated by a mtDNA distance of ca. 1% (Vázquez-Miranda, 2014). Of interest is the lack of differentiation in the large area occupied by the “northern” group, spanning from just north of the Vizcaíno Desert to Arizona, Sonora, Zacatecas, and Sinaloa. The nuclear gene tree (not shown) also supported two groups, albeit with relatively lower bootstrap frequencies, and an estimated time of divergence of 150,000 years (Vázquez-Miranda, 2014). A phylogenetic analysis of *Melanerpes* woodpeckers including two mtDNA genes, four nuDNA loci (1 Z-linked, 3 autosomal; Navarro-Sigüenza et al., 2017) also found north-south paraphyly. Thus, for the Gila woodpecker, previous results from the nuDNA loci and mtDNA suggest a shallow phylogeographic division in the vicinity of the Vizcaíno Desert.
- California gnatcatcher: The California gnatcatcher is distributed from southern California to the southern tip of Baja California Sur, and includes three described subspecies (see Mellink and Rea, 1994). Zink et al. (2000) surveyed mtDNA sequence variation from 64 individuals (1399 bp) throughout the range, finding no geographic divisions that were consistent with the limits of three described morphology-based subspecies (Atwood, 1991) or any other geographic units. Zink et al. (2013) found no geographic differentiation at a different mtDNA locus (ND2, 1041 bp), one Z-linked locus (529 bp), six autosomal introns (2331 bp) and one exon (506 bp). Similarly, in a phylogenetic analysis using one mtDNA gene and six nuDNA loci (2 Z-linked and 4 autosomal) no discernible geographic structure was recovered (Smith et al., 2018). Thus, for the California gnatcatcher, previous results from the nuDNA loci and the mtDNA gene tree were concordant and showed no genetic structure that coincides with subspecific designations of the California gnatcatcher or documented biogeographic breaks in Baja California (see U.S. Fish and Wildlife Service, 2011, 2016).
- LeConte’s thrasher: This species includes three subspecies, although the populations of *T*. *l*. *macmillanorum* are not considered valid by most authors (Sheppard, 2018). Zink et al. (1997) showed that populations north (*Toxostoma l*. *macmillanorum* plus *T*. *l*. *lecontei*) and south (*T*. *l*. *arenicola*) of the Vizcaíno desert were reciprocally monophyletic in mtDNA cytochrome *b* and ND6 sequences (619 bp). Vázquez-Miranda et al. (2017) sequenced the ND2 gene (1041 bp), seven Z-linked loci (4311 bp) and nine autosomal (6572 bp) loci and found the same split as reported by Zink et al. (1997) in both mitochondrial and nuclear gene trees. The ND2 gene tree showed a 2.08% sequence divergence between the two clades, five of seven Z-linked loci were fixed for one or two single nucleotide polymorphisms (SNPs), and one of nine autosomal loci showed a fixed difference (HMG, 2 SNPs). A minimum age of separation of 140,000 years was estimated for the two groups (Vázquez-Miranda et al., 2017). The predicted LGM distribution included two allopatric refugia (Vázquez-Miranda et al., 2017). Thus, for LeConte’s thrasher, previous results from the nuDNA loci and mtDNA were concordant and showed genetic structure supporting the divergence between subspecies (*T*. *l*. *lecontei* and *T*. *l*. *arenicola*), corresponding to the biogeographic break at the Vizcaíno Desert.

## 2. Methods

### 2.1. NGS data

Genotype by sequencing (GBS) data for Gila woodpecker, cactus wren, California gnatcatcher, and LeConte’s thrasher were generated and assembled using UNEAK pipeline in Tassel [v3.0.166] (Bradbury et al., 2007) at Cornell University. We received filtered SNP data from Cornell in HapMap format and converted each to VCF using TASSLE [v5.2.70] (Bradbury et al., 2007) and filtered to ensure that each biallelic SNP was present in ≥90% of individuals. Subsequently, we removed individuals from each dataset that possessed too much missing data for accurate downstream analyses: in cactus wren [COA36 (Coahuila), BCHVM121 (Cataviña), and CONACYT417 (Zacatecas)], in Gila woodpecker [two individuals from La Paz, BCHVM160 and BCHVM189], in California gnatcatcher [BCHVM118 and BCHVM223, Cataviña and Cabo Pulmo, respectively], and in LeConte’s thrasher [B16584 (Bahia de Santa Rosalillita) and BCHVM147 (Vizcaíno)], which brought the total amount of missing data per dataset to <10%.

For RADseq data in the California gnatcatcher, we generated libraries using previously published methods (Gamble et al., 2015). We first removed PCR duplicates using bbmap [v38.79] (Bushnell, 2014), then demultiplexed and filtered raw reads using STACKS [v2.53] (Catchen et al., 2013). We mapped reads to the Zebra finch genome (ENSEMBL, bTaeGut1_v1.p; Rhie et al., 2021) using bwa-mem2 [v2.1] (Li and Durbin, 2009; Vasimuddin et al., 2019) and assembled data using the ref_map.pl script in STACKS [v2.53], sampling one random SNP per locus. We further filtered these data such that each RADtag was biallelic, possessed a minimum read depth of 5, and was present in ≥90% of individuals. After assembling and filtering both GBS and RADseq datasets into comparable VCF files, we analyzed them in tandem for downstream analyses.

### 2.2 Population structure analyses, phylogenetic analyses, and ecological niche modeling

We examined the population genetic structure with each dataset using 3 methods; k-mean clustering using fastSTRUCTURE [v1.0.3] (Raj et al., 2014), Principal Components Analysis (PCA) using the R package, adegenet [v2.1.3] (Jombart, 2008), and phylogenies using IQ-TREE [v 2.0.3] (Nguyen et al., 2015). We converted all VCF files to the fastSTRUCTURE file format using PGD Spider [v2.1.1.5] (Lischer and Excoffier, 2012) and ran the program 5 times to explore K=1–5 for each dataset. Next, we used the chooseK.py script (part of the fastSTRUCTURE package) to identify a range of well-supported values of K and the R package, pophelper [v2.2.6] (Francis, 2017), to visualize each potential value of K. To further examine the genetic structure of these data, we used adegenet to conduct PCAs for each dataset. We converted each VCF to a FASTA using VCF-kit [v0.2.9] (https://github.com/AndersenLab/VCF-kit), filtered to include variable sites only using IQ-TREE, and constructed a maximum-likelihood phylogeny while correcting for ascertainment bias (GTR+ASC) inherent with SNP-based phylogeny construction (Leaché et al., 2015) to reduce phylogenomic distortion of branch lengths and topologies (Zink and Vázquez-Miranda, 2019) with statistical support from 1000 ultra-fast bootstrap (UFBoot) replicates (Hoang et al., 2018). Additionally, we estimated SNP species trees under an appropriate multispecies coalescent model using single value decomposition (Chifman and Kubatko, 2014) in SVDQUARTETS as part of PAUP* [v4.0a146] (D.Swofford, < http://people.sc.fsu.edu/~dswofford/paup_test/>), with 1000 non-parametric bootstrap replicates to assess branch support. Species tree methods allow us to estimate, incorporate and review conflicts between gene trees (Vázquez-Miranda and Barker, 2021). To visualize SNP gene-tree agreement and disagreement we estimated “cloudograms” of our species tree bootstrap replicates using PHANGORN (Schleip, 2011) in R [v3.5] (R Core Team, 2017). Lastly, we calculated *F*_*ST*_ across all SNPs and all taxa from the vcf files in VCFtools (Danecek et al., 2011) between clusters and clades to look at the distribution of levels of differentiation across Baja California. In case of no geographic structure, we used the accepted taxonomy for comparisons (Miller et al., 1957). We visualized levels of *F*_*ST*_ genomic differentiation using raincloud plots in R (code adapted from B. Warwick https://gist.githubusercontent.com/benmarwick/2a1bb0133ff568cbe28d/raw/fb53bd97121f7f9ce947837ef1a4c65a73bffb3f/geom_flat_violin.R and C. Scherer https://github.com/z3tt/TidyTuesday/blob/master/R/2020_31_PalmerPenguins.Rmd).

We constructed an ecological niche model (Peterson et al., 2011) for Gila woodpecker at the Last Glacial Maximum (LGM) and refer to similar models in Zink et al. (2013) for California gnatcatcher, Zink (2014) for cactus wren, and Vázquez-Miranda et al. (2017) for LeConte’s thrasher. Methods for the new Gila woodpecker niche model (Higmans et al., 2005) followed those of Zink (2014). It is likely that divergence of some taxa in this study preceded the LGM. However, we suggest that it is relevant to use niche models to identify potential LGM refugia that could have allowed the persistence of phylogroups that diverged prior to the LGM. That is, if there were no allopatric refugia at the LGM, prior differentiation would have disappeared because of intermingling of once-distinct groups.

## 3. Results

### 3.1 Data description

Basic single locus data for cactus wren and Gila woodpecker are given in Table 1. The final, filtered genome-wide SNP data for each species were as follows: the cactus wren GBS dataset contained 14,874 biallelic SNPs with 5.7% missing data; the Gila woodpecker GBS dataset contained 25,512 biallelic SNPs with 4.3% missing data; the California gnatcatcher GBS dataset contained 33,806 biallelic SNPs with 3.2% missing data; the California gnatcatcher RADseq dataset contained 8,734 biallelic SNPs with 2.1% missing data; the LeConte’s thrasher GBS dataset contained 24,833 biallelic SNPs with 2.8% missing data.

**Table 1:** Locus by locus summary of sample sizes and genetic results for cactus wren and Gila woodpecker (see Zink et al., 2013; 2016) for data from California gnatcatcher and Vázquez-Miranda et al. (2017) for LeConte’s thrasher).

| Locus (base pairs) | South (n,<br>number<br>haplotypes, pi) | North (n,<br>number<br>haplotypes, pi) | $F_{ST}$ |
| --- | --- | --- | --- |
| **Cactus wren** |  |  |  |
| ND2 (1041) | 10,7,0.0040 | 9,7,0.0037 | 0.926*** |
| ACL (479) | 10,9,0.0056 | 6,5,0.0061 | 0.385 |
| TGFB2 (592) | 10,4,0.0027 | 10,4,0.0026 | 0.377* |
| AETC (376) | 10,3,0.002 | 10,1,0.0 | 0.476** |
| MPP (378) | 14,4,0.0028 | 10,3,0.0016 | 0.565** |
| GAPDH (340) | 10,2,0.0006 | 10,4,0.0026 | 0.302* |
| ACON1 (1027) | 14,4,0.0009 | 10,3,0.0007 | 0.628** |
|  |  | 12,5,0.225 |  |
| **Gila woodpecker** |  |  |  |
| ND2 (1041) | 20,6, 0.0010 | 41,15,0.0014 | 0.75*** |
| FIB7 (820) | 28,2,0.0002 | 57,6,0.0010 | 0.055* |
| ACO1 (1044) | 28,4,0.0009 | 58,9,0.0007 | 0.029 |
| TGFB2 (585) | 3,0,0 | 4,0,0 | 0.0 |
| GAPDH (461) | 8,1,0 | 10,3,0.0016 | 0.267 |

### 3.2 Population genetic structure

Our fastSTRUCTURE analyses suggested the best value (or range of values) of K using the chooseK.py script for each dataset are presented in Figure 2 and are as follows: for the cactus wren GBS dataset, K = 3; for the Gila woodpecker GBS dataset, K = 1–2; for the California gnatcatcher GBS, K = 1–2; for the California gnatcatcher RADseq dataset, K = 2; for the LeConte’s thrasher dataset GBS, K = 2–3. We qualitatively corroborated the likely values of K using PCAs for each dataset, especially those where a range of K values was specified previously. For each dataset, we identified/confirmed the most likely value of K as follows: cactus wren GBS (K = 3), Gila woodpecker GBS (K = 2; no outgroup present in this dataset), California gnatcatcher GBS (K = 2), California gnatcatcher RADseq (K = 2), and LeConte’s thrasher GBS (K = 3) (Fig. 2). Details of genetic clustering information is available for each species/dataset in Supplemental Figures 4 – 8.

**Figure 2:**
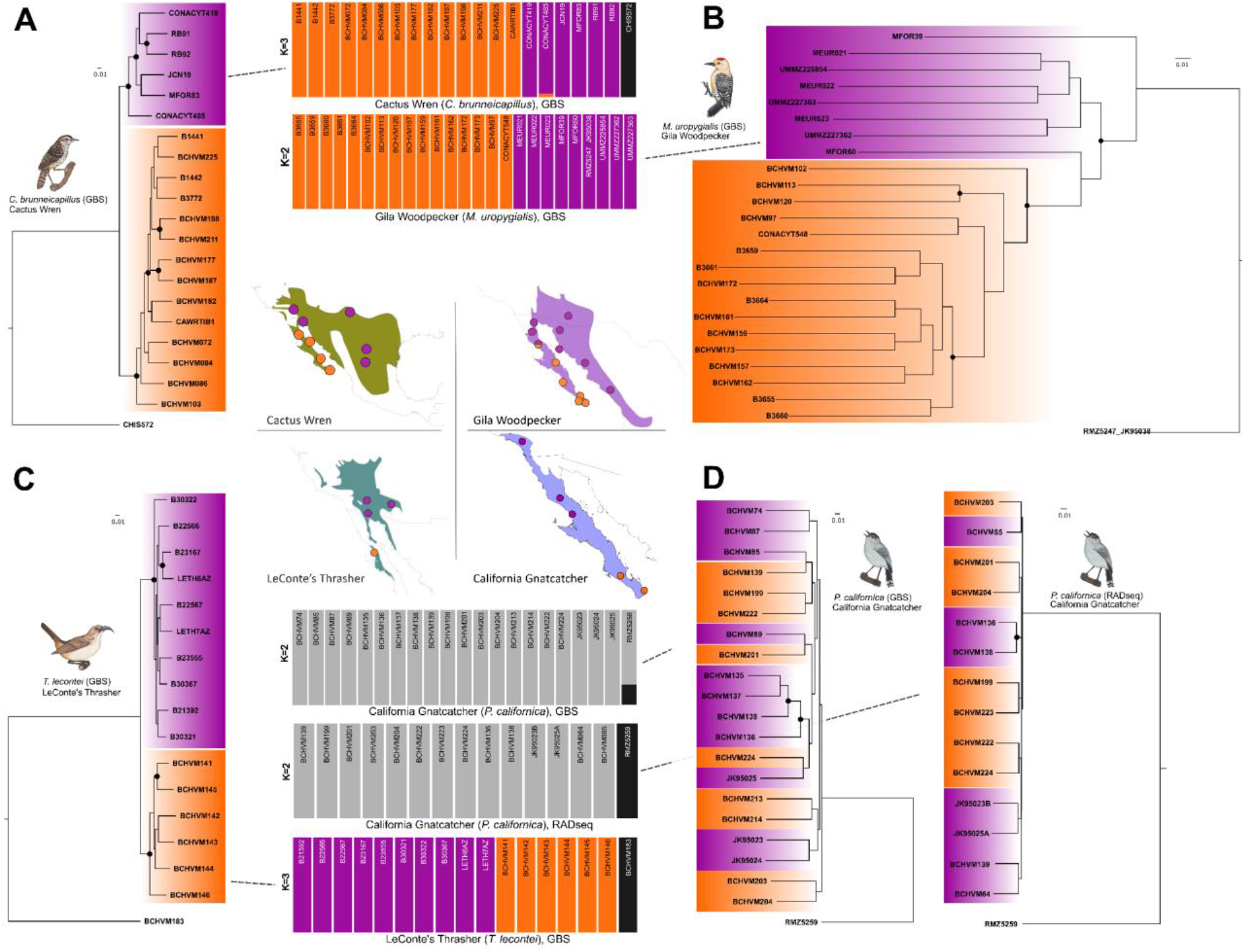
Sampling and phylogenetic structure of the four focal species of Baja birds: (A) Cactus Wren (*C*. *brunneicapillus*), (B) Gila Woodpecker (*M*. *uropygialis*), (C) LeConte’s Thrasher (*T*. *lecontei*), and (D) the California Gnatcatcher (*P*. *californica*, GBS—left/RADseq—right)). Map dots show general localities. Color coordination between maps, phylogenetic trees, and structure plots indicate diagnosable geographic population structure that coincides with north (darker/purple) or south (lighter/orange) of the Vizcaíno desert, Baja California. Gray in structure plots indicate a lack of diagnosable genetic clustering—outgroups are coded charcoal grey/black, if available. Black circles on phylogenies indicate well-supported nodes (UFBoot values ≥95). All specimens from Bell Museum, University of Minnesota. Bird illustrations by A. Gardner.

The ML phylogeny for each dataset possessed only a small number of well-supported nodes (Fig. 2). However, these nodes we generally located deeper in the phylogeny indicating that, if it were there, we would likely be able to detect deeper phylogenetic structure within these focal species. Not all species have congruent distributions (Fig. 2) although samples for most species overlap.

- Cactus wren: Samples fall into two groups, California and Mexican mainland, and samples from Cataviña southward; the samples from Cataviña sort out paraphyletically at the base of the Baja cluster (Fig. 2).
- Gila woodpecker: We resolved a complex structure within our samples. Indeed, we resolved one poorly-resolved group of southern Baja samples with a clade of northern Baja/mainland samples nested deep within (Fig. 2). These results correspond with the predicted LGM distribution (Fig. S4) suggested two allopatric refugia.
- California gnatcatcher: The two datasets for California gnatcatcher failed to recover any notable geographic structure (Fig. 2).
- LeConte’s thrasher: There was a distinct division of populations from the Vizcaíno Desert and those to the north (Fig. 2).

Species-tree analyses (Fig. S5 - 9) were consistent with ML phylogenies (Fig. 2) but showed 100% bootstrap support for all nodes and all taxa, except for Gila woodpeckers. LeConte’s thrashers and cactus wrens were sorted north and south, Gila woodpeckers in the north were paraphyletic to those in the south, and California gnatcatchers were unsorted. Consistent with little to no gene-tree conflict, all cloudograms showed the same topology as ML analyses with no “fuzziness” representing alternate topologies. We detected gene-tree conflict in Gila woodpeckers evidenced by its cloudogram’s fuzziness, although northern individuals tended to cluster with themselves separate from southern individuals. We found a gradient of genomic divergence across all taxa. Following the population structure and phylogenetic results, LeConte’s thrashers and cactus wrens showed *F*_*ST*_ values closer to 1.0 with many loci fixed between north and south, followed by Gila woodpeckers with values >0.9. In contrast, *F*_*ST*_ values for California gnatcatchers (GBS and RADseq) were closer to 0.0, with only a handful of SNPs passing the 0.75 landmark. The average *F*_*ST*_ -values for those genome-wide SNPs were in descending order: LeConte’s thrashers 0.20 (−0.08-0.48 CI); cactus wrens 0.18 (−0.08-0.44 CI); Gila woodpeckers 0.08 (−0.06-0.22 CI); California gnatcatchers RADseq 0.05 (−0.05-0.15 CI) and GBS 0.04 (−0.04-0.11 CI).

## 4. Discussion

### 4.1 Summary of phylogenomic results with disparate data

- Cactus wren: Previous results from the nuDNA loci and mtDNA were concordant and showed a distinct phylogeographic split at the implied location of the ancient mid-peninsular seaway (latitude 27°–30°N). Here, our GBS data are consistent with this hypothesis.
- Gila woodpecker: Previous results from the nuclear nuDNA loci and mtDNA suggested relatively weak phylogeographic structuring. Our GBS data suggest that northern Baja and mainland populations are nested within populations from southern Baja (Fig. 2). The division between this monophyletic northern group also lies at the implied location of the ancient mid-peninsular seaway (latitude 27°–30°N). The scenario of a northern clade nested within southern samples is most consistent with a northern dispersal/vicariance from a original southern Baja population.
- California gnatcatcher: Previous results from the nuDNA loci and mtDNA were concordant and showed no genetic structure that coincides with subspecific designations of the California gnatcatcher. Here, our GBS and RADseq datasets both support a lack of genetic structure upon which subspecies could be based.
- LeConte’s thrasher: Previous results from the nuDNA loci and mtDNA were concordant and showed genetic structure supporting the divergence between populations from the mainland and southern Baja (nominal subspecies: *T*. *l*. *lecontei* and *T*. *l*. *arenicola*). Here, our GBS data support this phylogeographic division.

Given concerns about the exclusive use of mtDNA, or small samples of nuclear loci (Teske et al., 2018; Drovetski et al., 2015) it is useful to investigate large samples of loci from next-generation sequencing methods to test earlier results from less comprehensive methods. Even if on average nuclear loci coalesce too slowly to capture very recent population fission events, sampling thousands of loci provides the opportunity to discover loci with more rapid than average coalescence times. In the species we examined, GBS and RADseq data show consistency with results from mtDNA and single nuclear gene loci, all of which reveal concordant phylogeographic gaps in LeConte’s Thrasher, cactus Wren, and Gila Woodpecker, centered near the Vizcaíno Desert, and the absence of genetic divisions within the California gnatcatcher. The weak separation of samples of Gila woodpecker north and south of the Vizcaíno Desert is consistent with the low level of mtDNA separation, and likely a result of the differences in coalescence times of the two sets of markers (Zink and Barrowclough, 2008). Although our genome-wide sample of loci from the GBS technique nonetheless represents a small part of the genome, the congruence across molecular markers suggests that the recent phylogeographic history of these species can be reliably inferred and that single locus biases or incompatibilities between markers are not obscuring interpretations of their differentiation (Hill 2015).

### 4.2 Taxonomy, phylogeography and conservation implications

Correct identification of units of biodiversity is imperative for defining sensible conservation plans (Ely et al., 2017). With limited resources for species listed as threatened or endangered, it is important that units identified for protection are evolutionarily distinct (Moritz, 1994). Although most intraspecific divisions, typically subspecies, were based on aspects of the external morphology, such as plumage coloration and size and shape, data derived from molecular methods have become more accepted proxies for lineage independence owing to the fact that their genetic bases and evolutionary transitions are understood. That is not to say that discrete, concordant morphological differences should not be recognized taxonomically at some level, such as for the spotted owl (*Strix occidentalis*), where morphology and genetics agree on the validity of three species-level taxa (Barrowclough et al., 2005, 2011). In contrast, a large number of avian subspecies were not supported by examination of the geography of mtDNA gene trees (Zink, 2004; Phillimore and Owens, 2005). It is possible that significant geographic and adaptive variation might be influenced by just a few loci and therefore previous techniques could have excluded the pertinent loci (Toews et al., 2015). Thus, whether subspecies qualify as evolutionarily independent taxa is unclear, and it is possible that their evolutionary validity is supported by too few genes to be found by surveys of mtDNA or a few nuclear loci, but might be overcome by genome-wide SNP data such as those presented here.

The current species-level and subspecies-level taxonomy does not capture the evolutionary diversity in the species examined here. LeConte’s thrasher includes two distinct evolutionarily groups that are consistent with currently recognized subspecies. In the cactus wren, seven subspecies (Miller et al., 1957; Rea and Weaver, 1990) are represented in the two distinct evolutionary lineages (north: *C*. *b*. *couesi*, *C*. *b*. *bryanti*, *C*. *b*. *seri* [Isla Tiburon], *C*. *b*. *brunneicapillus*, *C*. *b*. *guttatus*; south: *C*. *b*. *affinis and C*. *b*. *purus*). The phylogeographic gap falls more or less in the middle of the range of *C*. *b*. *purus* (29^°^ to 35^°^). For the Gila woodpecker, our potential southern Baja California clade would be consistent with the distribution of *M*. *u*. *brewsteri* (28° south to Cabo San Lucas), whereas the northern clade contains individuals representing five subspecies (*M*. *u*. *uropygialis*, *M*. *u*. *tiburonensis*, *M*. *u*. *albescens*, *M*. *u*. *fuscescens*, *M*. *u*. *cardonensis*) (Miller et al., 1957). Thus, the cactus wren and LeConte’s thrasher could each be considered as two species divided at the Vizcaíno Desert, which would acknowledge their concordant evolutionary histories with each other and the other co-distributed taxa (Alvarez-Castanedaand Murphy, 2014; Dolby et al., 2015). Certainly, taxonomic revision will depend on the method and philosophy of species delineation (Cicero et al., 2021). For the Gila woodpecker, it is possible that populations north and south of the Vizcaíno Desert could be considered separate species even though we did not find evidence of reciprocal monophyly. However, it would be prudent to generate additional data to identify the root of the tree more effectively before recommending specific taxonomic changes.

Originally thought to be conspecific with the black-tailed gnatcatcher (*P*. *melanura*) the California gnatcatcher was shown to be morphologically and genetically distinct (Atwood, 1991; Zink and Blackwell-Rago, 1998). *Polioptila c*. *californica*, known as the coastal California gnatcatcher, as delimited by Atwood (1991) was listed in 1993 (USFWS, 1993) as threatened under the US Endangered Species Act (ESA). Since this description, there has been a complex and controversial history involving the subspecific taxonomy of the California gnatcatcher (for review see: Skalski et al., 2008; McCormack and Maley, 2015; Smith et al., 2018; Vandergast et al., 2019; Zink et al., 2000, 2013, 2016). In brief, over the past ca. two decades the status of *P*. *c*. *californica* has been called into question using genetic and morphological data with varying statistical power. Here, we show using two different genome-wide SNP datasets that the California gnatcatcher is a single panmictic population. In particular, the distribution of *F*_*ST*_ values (Fig. 3b) shows that, across its range, the California gnatcatcher is less differentiated overall than other co-distributed taxa (i.e. Gila woodpecker, cactus wren, or LeConte’s thrasher). As stated by the Sesame Street crew in their famous song, “one thing is not like the three others”. Given that current taxonomy rates the latter three taxa as subspecies, it would appear that given the genetic data derived from GBS and RADseq analyses that no taxonomic recognition is warranted for the gnatcatchers.

**Figure 3.**
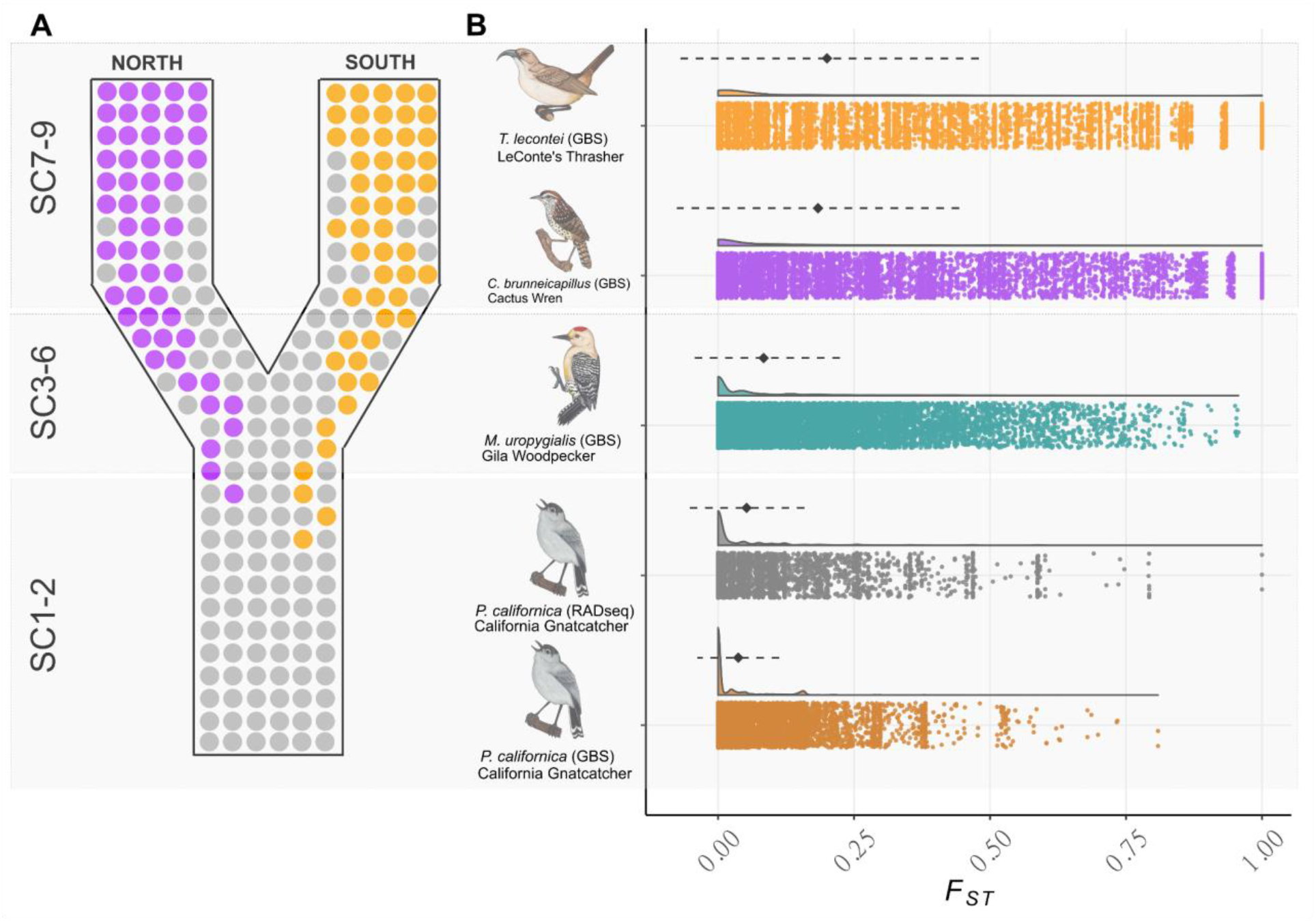
The nature of lineage divergence and gradients of genomic differentiation. (A). The time course of lineage differentiation (de Queiroz, 2007 and corresponding species criteria [SCs 1-9]; and Leliaert et al., 2014). (B). Gradient distributions of *F*_*ST*_ values from genome-wide SNP comparisons between clades for sorted taxa, and taxonomy for unsorted taxa. Diamonds and dotted lines represent mean *F*_*ST*_ values and confidence intervals (CI), respectively. Bird illustrations by A. Gardner.

### 4.3 Comparative phylogeography

Strong barriers to dispersal ought to create congruent patterns of divergence within species whose ranges span them irrespective of different habitat and life history characteristics. However, several factors can mitigate common responses to barriers, including dispersal abilities, effective population size and species ecology. We also surveyed small samples of ten additional species for mitochondrial ND2 (1041 base pairs) and found five species that appeared undifferentiated (*F*_*ST*_ not significant) across the Vizcaíno desert: (California quail, *Callipepla californica*; Bewick’s wren, *Thyromanes bewickii*; black-throated sparrow, *Amphispiza bilineata*; spotted towhee, *Pipilo maculatus*; house finch, *Haemorhous mexicanus*), and four species that were differentiated (verdin, *Auriparus flaviceps*; California towhee, *Melozone crissalis*; ladder-backed woodpecker, *Picoides scalaris*; ash-throated flycatcher, *Myiarchus cinerascens*) (Table S2). Thus, in total there are six undifferentiated (those above plus California gnatcatcher) and eight differentiated species (those mentioned above plus cactus wren, LeConte’s thrasher, Gila woodpecker, and savannah sparrow (Zink et al., 2005; Benham and Cheviron, 2019). Thus, it appears that if there was mid-peninsular seaway that led to splits north and south of the Vizcaíno Desert, it was (1) of insufficient duration to affect all species that were co-distributed on either side at the time it was present, (2) it was not a barrier to some species, or (3) some species rapidly recolonized either the north or southern parts of the historic range once the seaway resided. Zink et al. (2000) suggested that the California gnatcatcher was a relatively recent arrival to the coastal sage scrub north of the Vizcaíno Desert, having relatively recently dispersed northward from a southern Baja California LGM refugium (insufficient samples preclude a similar analysis with GBS or RADseq data). Multilocus data also show evidences of post-LGM demographic expansions on either side of the Vizcaíno Desert (Zink et al., 2013). These, or other alternatives, require more and larger samples to distinguish between them. Lastly, our data (Table S2) on small samples from the Sierra de la Laguna corroborate previous studies (Honey-Escandón et al., 2008; Walstrom et al., 2012; Friis et al., 2016). Thus, there are at least two major biogeographic breaks in the avifauna of Baja California.

### 4.4 Stages in the process of lineage divergence

The time course of lineage divergence passes through successive stages of polyphyly, paraphyly, and—eventually—reciprocal monophyly (de Queiroz, 2007; Leliaert et al., 2014). This process can take a long time depending on effective population sizes. Our comparative phylogenomic study shows a gradient of differentiation among co-distributed avian taxa across Baja California. Thrashers and wrens have reached the point of reciprocal monophyly, woodpeckers are intermediate and paraphyletic, and gnatcatchers are polyphyletic, and either at the start of the process of lineage divergence or at an equilibrium maintained by gene flow. The ultimate fate of these lineages will depend on many factors, including the strength of ecological barriers, dispersal ability and effective population sizes. The stages of lineage divergence are recognized taxonomically in different ways depending on which subspecies and species concepts are used. Vázquez-Miranda et al. (2017) suggested that the two groups of LeConte’s thrashers have been separated for at least 140,000 years and given the short geographic distance separating the two clades, there has been ample time for introgression, of which there is no evidence. Thus, this species, as well as the cactus wren, have achieved the status of phylogenetic species, whereas their status of biological species is determined by a vote of the North American Classification and Nomenclature Committee. Gila woodpeckers are at an evolutionary crossroads: they have diverged north-south yet they still exhibit incomplete lineage sorting and/or gene flow. Whether the California gnatcatcher will eventually reach the stage of multiple lineages as found in the LeConte’s thrasher and cactus wren is unknown, although there appear to be no biogeographic barriers that would impede gene flow (other than geographic distance). These comparisons illustrate the relationship between lineage divergence and speciation as an extended process (Avise and Walker, 1998; Shaw and Mullen, 2014; Stankowski and Ravinet, 2021).

### 4.5 Preserving areas of endemism in conservation planning

Taxonomic resolution leads to recommendations for conservation planning. The Sierra de la Laguna is recognized as the “Sierra La Laguna Biosphere Reserve” by the United Nations (UNESCO, https://en.unesco.org/biosphere/lac/sierra-la-laguna) in recognition of its unique biological diversity. Our results corroborate the distinctiveness of species in that area. At the community level, our results are consistent with many other studies (Riddle et al., 2000; Garrick et al., 2009) that reveal a significantly differentiated biota south of the Vizcaíno Desert, and in the event entire reserves were to be set aside, genetic results could guide their delimitation. For example, the El Vizcaíno Biosphere Reserve (http://www.parkswatch.org/parkprofile.php?l=eng&country=mex&park=vibr&page=phy) includes the southern clade of the LeConte’s thrasher (*Toxostoma lecontei arenicola*) as well as other species mentioned in this study. Such reserves are an excellent example of preserving broad patterns of biodiversity.

### 4.6 Conclusion

In sum, we examined phylogeographic patterns in multiple species distributed across Baja California. We show that previous work using mitochondrial markers and nuclear markers are largely concordant with similar analyses conducted with GBS and RADseq data for four bird species found along the Baja California peninsula and adjacent mainland (cactus wren, Gila woodpecker, California gnatcatcher, and LeConte’s thrasher). Three of four species show a mostly concordant historical division at or near the Vizcaíno Desert in north-central Baja California. For 12 other species, we found a roughly even number of differentiated and undifferentiated species across the Vizcaíno Desert—with no obvious explanations for the difference. Thus, these data will help generate future taxonomic and biogeographic hypotheses for how flora and fauna evolved across the Baja California peninsula in Mexico and inform conservation planning.

## Acknowledgments

The authors thank A. Gardner for providing illustrations of the focal taxa in this study (Figures 2, 3). K. Barker, K. Burns and A. Navarro-Sigüenza provided locality information. The University of Washington Burke Museum, The Field Museum of Natural History, the San Diego Natural History Museum, University of California (Berkeley) Museum of Vertebrate Zoology, Museo de Zoología “Alfonso L. Herrera” (MZFC), Facultad de Ciencias, UNAM, San Diego State University, University of Michigan Museum of Zoology, Louisiana State University Museum of Natural Science, provided tissue samples. Support was received from several University of Minnesota sources: the Dayton-Wilkie fund at the Bell Museum of Natural History, an International Graduate Student Grant from the Graduate School, two Huempfner Fellowships from the College of Biological Sciences, and one Anderson Fellowship. Funding was also received from the Chapman Fund of the American Museum of Natural History, the American Ornithologists’ Union, the Consejo Nacional de Ciencia y Tecnología of Mexico (CONACyT), and the Universidad Nacional Autónoma de México (UNAM DGAPA-PAPIIT204220). Bioinformatic analyses were partially performed at the University of Nebraska-Lincoln (UNL) Holland Computer Center (HCC). The authors declare that they have no known competing financial interests or personal relationships that could have appeared to influence the work reported in this paper.

## Author contributions

H.V-M. designed study, conducted fieldwork, performed laboratory work, analyzed data, drafted an early stage of the manuscript and secured funding. R.M.Z. conceptualized study, performed genetic analyses, conducted fieldwork, and wrote the manuscript. B.J.P. conducted final genome-wide SNP analyses, wrote associated sections, and helped write the manuscript. All authors read and approved the manuscript for submission.

## Data Availability

All genome-wide SNP data used to generate the analyses and figures presented in this manuscript is openly available after publication on Figshare at https://doi.org/10.6084/m9.figshare.14368301. In addition, raw reads generated for the *P*. *californica* (California Gnatcatcher) RADseq analyses are available on the NCBI SRA under BioProject PRJNA719408. Single marker mtDNA and nuclear loci used in this study are available on GenBank under accessions MZ476275 - MZ476525. For figures, the following specimen labels are from these museums: Bell Museum, University of Minnesota (BCHVM, LETH, CAWR, MEUR, RMZ); Louisiana State University Museum of Natural Science (B); University of Michigan Museum of Zoology (UMMZ); Field Museum of Natural History (FMNH, BRUN); Burke Museum University of Washington (JK, RB); Museo de Zoología “Alfonso L. Herrera”, Universidad Nacional Autónoma de México (CONACYT, CAWR, SAR, CSW, VGR, EAD, MFOR).

