## Supplementary material for "Comparative phylogenomic patterns in the Baja California avifauna, their conservation implications, and the stages in lineage divergence": Table S1

**Table S1.** Localities for specimens (row correspond to main text Figure 1).

| General Locality | Longitude | Latitude |
|---|---|---|
| 1. San Diego | -116.83 | 32.82 |
| 2. Laguna Salada | -115.70 | 32.11 |
| 3. Ensenada | -116.60 | 31.87 |
| 4. Sierra Las Tinajas | -115.43 | 31.83 |
| 5. San Felipe | -114.96 | 30.59 |
| 6. Misión San Fernando | -115.20 | 29.98 |
| 7. Cataviña | -114.91 | 29.67 |
| 8. Isla Tiburón | -112.40 | 28.90 |
| 9. Santa Rosalillita | -114.19 | 28.70 |
| 10. Vizcaíno (Bahía Tortugas) | -113.89 | 27.45 |
| 11. San Ignacio | -112.90 | 27.28 |
| 12. San Javier (Rancho Monte Alto) | -111.62 | 25.93 |
| 13. Ciudad Insurgentes | -111.78 | 25.26 |
| 14. El Cien | -110.98 | 24.37 |
| 15. La Paz | -110.40 | 24.29 |
| 16. Rancho La Ballena | -109.97 | 23.74 |
| 17. Sierra de la Laguna | -109.98 | 23.65 |
| 18. San Dionisio | -109.87 | 23.56 |
| 19. Todos Santos | -110.23 | 23.45 |
| 20. Miraflores | -109.78 | 23.37 |
| 21. Santa Anita | -109.70 | 23.18 |
| 22. Cabo Pulmo | -109.51 | 23.56 |

**Table S2.** ND2 results for samples of selected species south and north of the Vizcaíno desert (Vázquez-Miranda et al., unpubl. data). Species denoted with * involve comparisons of samples from the Sierra de la Laguna and elsewhere. Calculations performed with DnaSP (Rozas et al., 2017).

| **Locus (base pairs)** | **South (n, number haplotypes, pi)** | **North (n, number haplotypes, pi)** | $\boldsymbol{F_{ST}}$ |
|---|---|---|---|
| **Verdin, *Auriparus flaviceps*** | | | |
| ND2 (1041) | 5,2,0.0008 | 11,8,0.0025 | 0.969*** |
| **California quail, *Callipepla californica*** | | | |
| ND2 (1041) | 5,4,0.0021 | 4,4,0.0014 | 0.245 |
| **California towhee, *Melozone crissalis*** | | | |
| ND2 (1041) | 5,4,0.0035 | 11,5,0.0013 | 0.235* |
| **Bushtit*, *Psaltriparus minimus*** | | | |
| ND2 (1041) | 4,1,0.0 | 5,3,0.0015 | 0.875* |
| **Bewicks, wren, *Thyromanes bewickii*** | | | |
| ND2 (1041) | 2,2,0.001 | 5,4,0.0015 | 0.0 |
| **Ladder-backed woodpecker, *Picoides scalaris*** | | | |
| ND2 (1041) | 8,3,0.0018 | 5,4,0.0005 | 0.823* |
| **Black-throated sparrow, *Amphispiza bilineata*** | | | |
| ND2 (1041) | 10,5,0.0019 | 7,6,0.0047 | 0.041 |

| | | | |
|---|---|---|---|
| **Acorn woodpecker*, *Melanerpes formicivorus*** | | | |
| ND2 (1041) | 4,1,0.0 | 5,3,0.0008 | 0.946* |
| **Ash-throated flycatcher, *Myiarchus cinerascens*** | | | |
| ND2 (1041) | 5,2,0.0004 | 6,3,0.0008 | 0.627* |
| **San Lucas robin*, *Turdus confinus*** | | | |
| ND2 (1041) | 5,2,0.0004 | 3,3,0.0019 | 0.87* |
| **White-breasted nuthatch*, *Sitta carolinensis*** | | | |
| ND2 (1041) | 4,1,0.0 | 8,4,0.0014 | 0.99* |
| **Baird's (dark-eyed) junco*, *Junco bairdi*** | | | |
| ND2 (1041) | 8,1,0.0 | 12,2,0.0074 | 0.99*** |
| **Spotted towhee, *Pipilo maculatus*** | | | |
| ND2 (1041) | 5,1,0 | 5,5,0.006 | 0.51 |
| **Savannah sparrow, *Passerculus sandwichensis*** | | | |
| ND2 (1041) | 35,6,0.0008 | 14,8,0.012 | 0.413** |
| **House finch, *Haemorhous mexicanus*** | | | |
| ND2 (1041) | 5,4,0.0015 | 31,31,0.0033 | 0.032 |

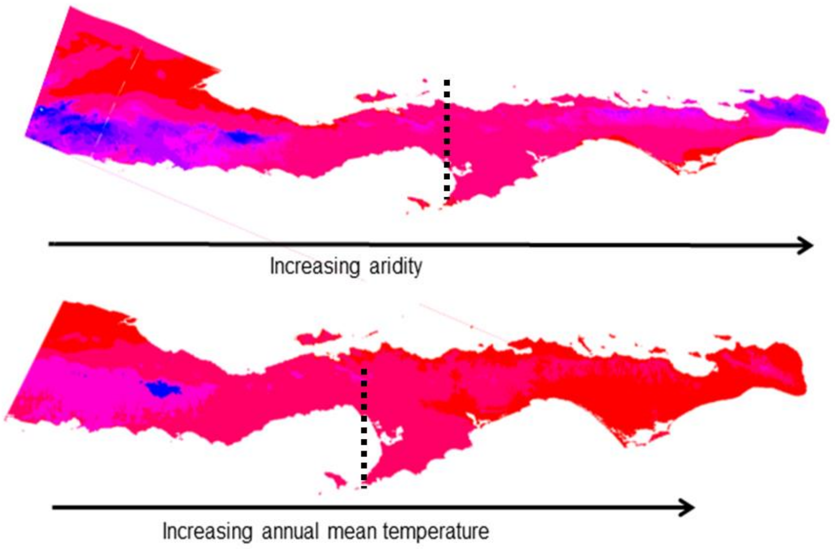


**Figure S1:** Patterns of annual temperature and aridity in Baja California (from Worldclim, Hijmans et al. (2005) WorldClim, version 1.3. University of California, Berkeley, CA). Available at: https://biogeo.berkeley.edu/worldclim/worldclim.htm.). Dotted line at approximate location of Vizcaíno Desert and phylogeographic breaks.

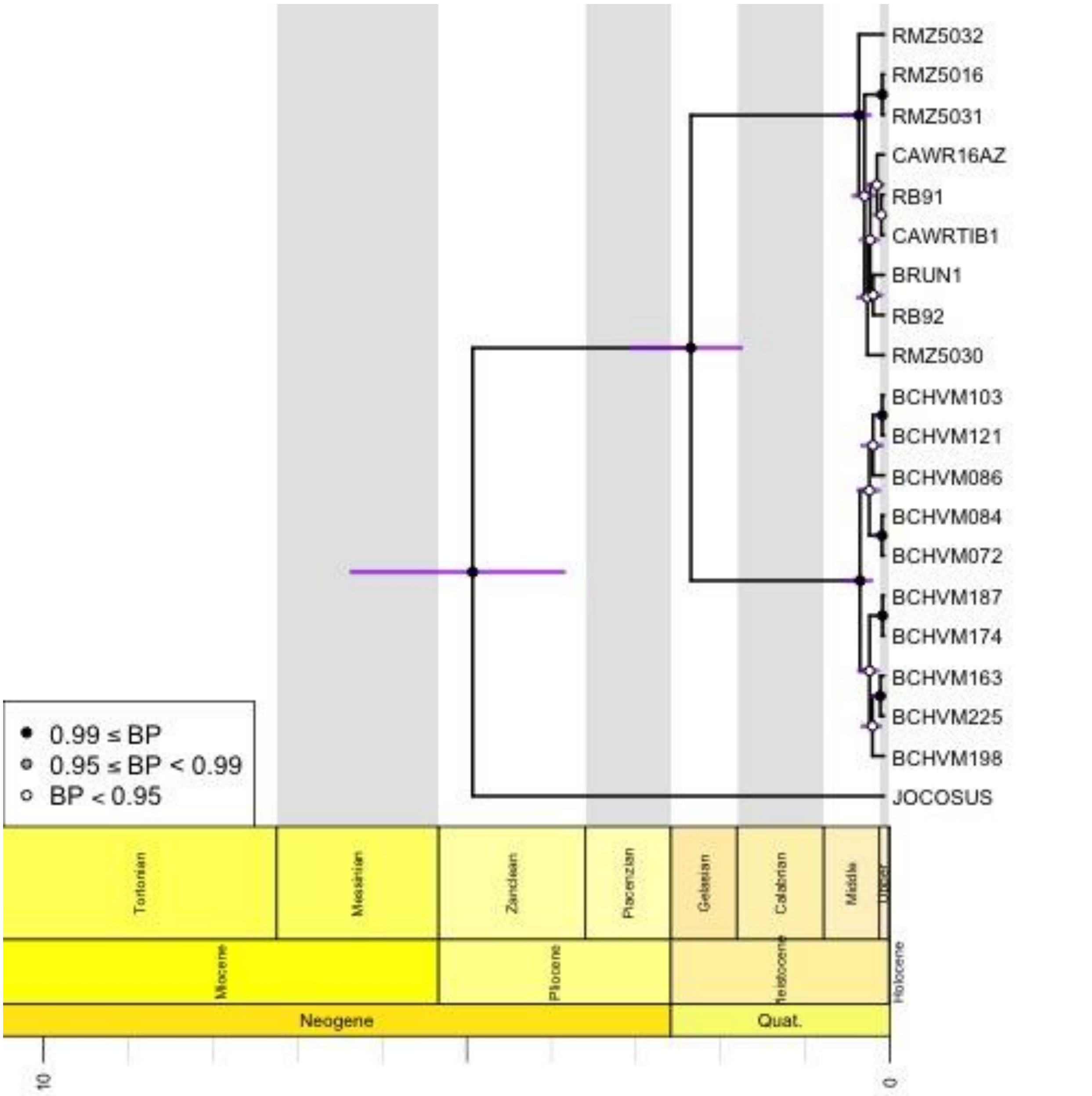


**Figure S2:** Mitochondrial DNA strict-clock chronograms based on ND2 from BEAST (Drummond et al., 2012). Tree rooted with *Campylorhynchus jocosus* (JOCOSUS; Genbank accession MZ380538; Vázquez-Miranda and Barker, 2021). Bars on chronograms represent node height Highest Posterior Density intervals (age). Node color values denote posterior probability support (see legend). Scale axes are in million years before present. See Data Availability section for explanation of specimen labels.

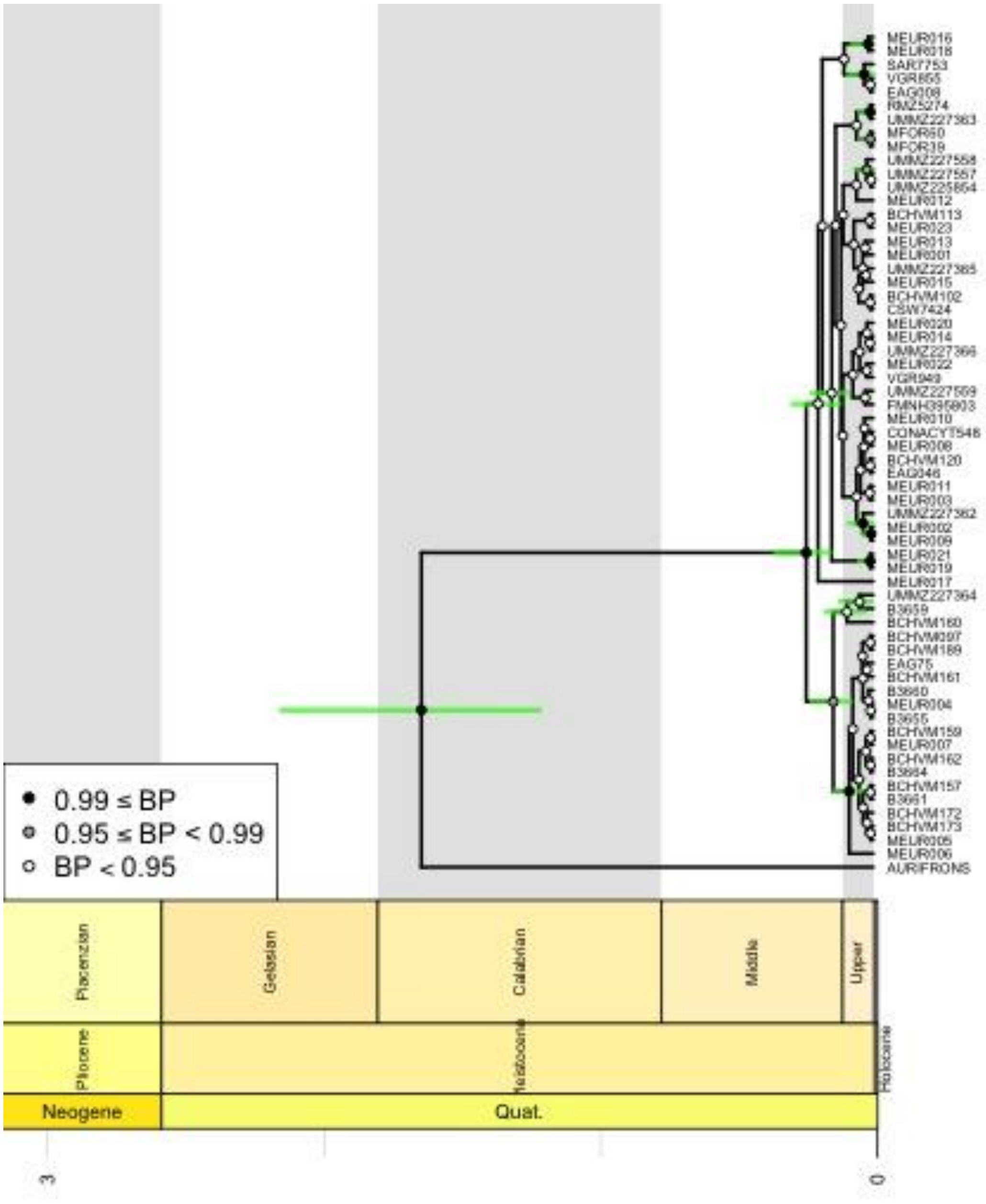


**Figure S3:** Mitochondrial DNA strict-clock chronograms based on ND2 from BEAST (Drummond et al. 2012). Tree rooted with *Melanerpes aurifrons* (AURIFRONS; Genbank accession KY288596; Navarro-Sigüenza et al. 2017). Bars on chronograms represent node height Highest Posterior Density intervals (age). Node color values denote posterior probability support (see legend). Scale axes are in million years before present. See Data Availability section for explanation of specimen labels.

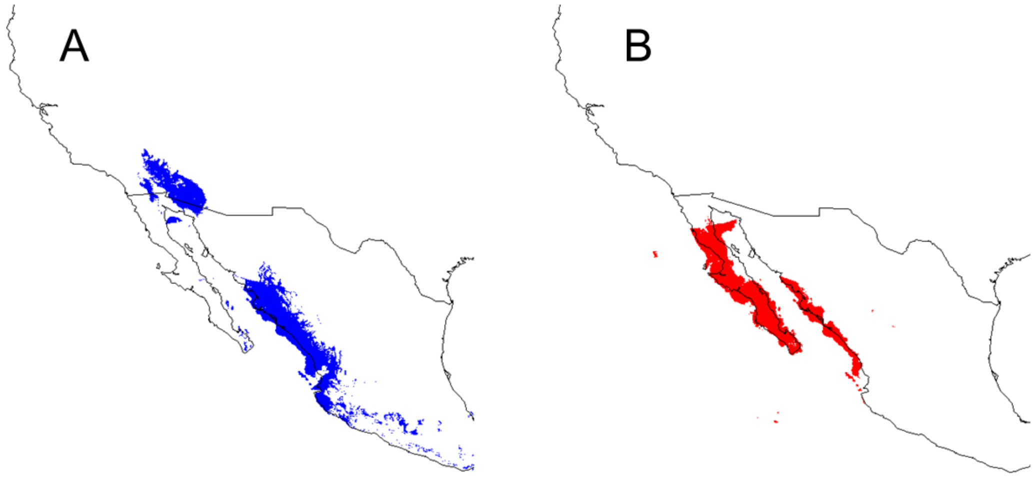


**Figure S4:** Ecological niche models showing predicted distributions of Melanerpes uropygialis at the Last Glacial Maximum. A. model built excluding localities from Baja California, B. model using only localities from Baja California.

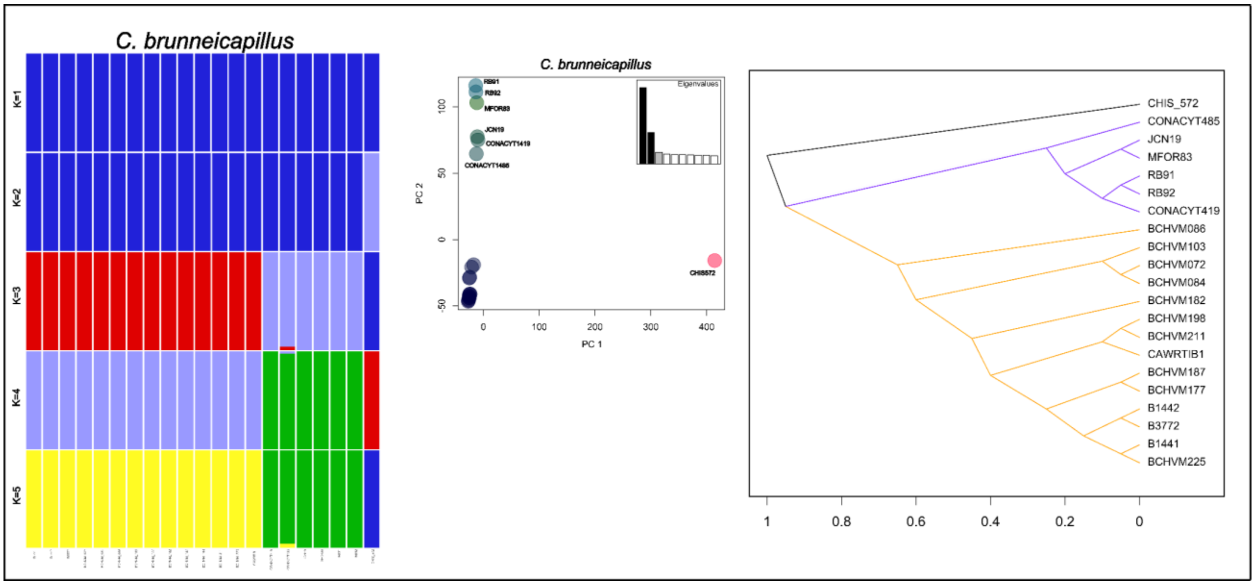


**Fig. S5.** Population clustering information for Cactus wren (GBS). Structure plots (K=1-5), PCA, and SNP cloudogram, respectively. PCA points labeled as individuals, when applicable, remaining unlabeled samples cluster too closely to be readily distinguishable. All voucher tissue samples in the Bell Museum, University of Minnesota. See methods for additional details.

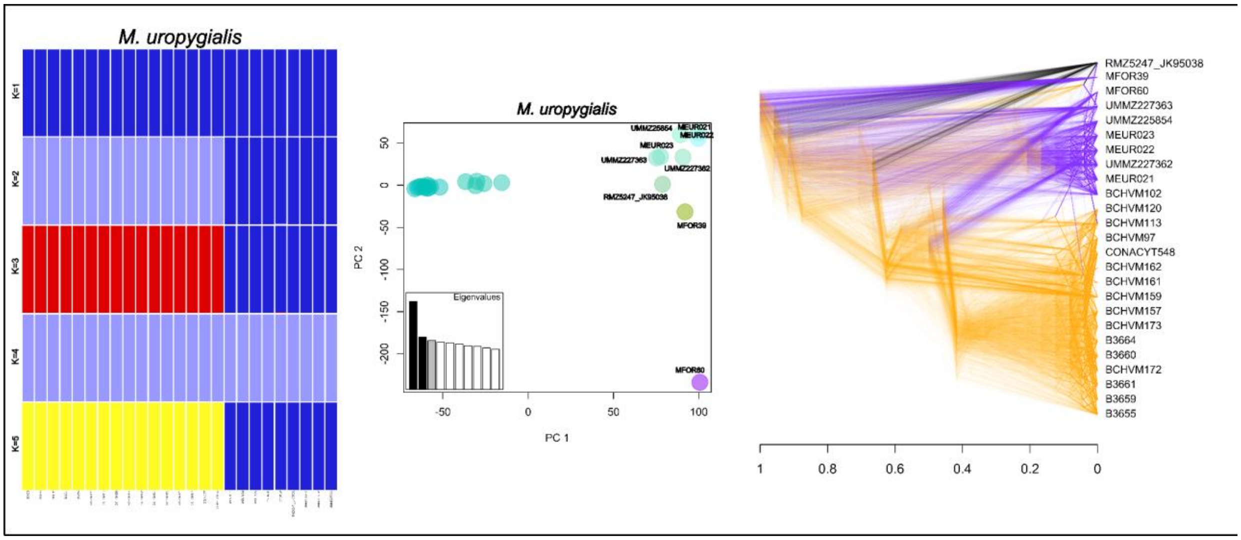


**Fig. S6.** Population clustering information for Gila Woodpecker (GBS). Structure plots (K=1-5), PCA, and SNP cloudogram, respectively. PCA points labeled as individuals, when applicable, remaining unlabeled samples cluster too closely to be readily distinguishable. See methods for additional details.

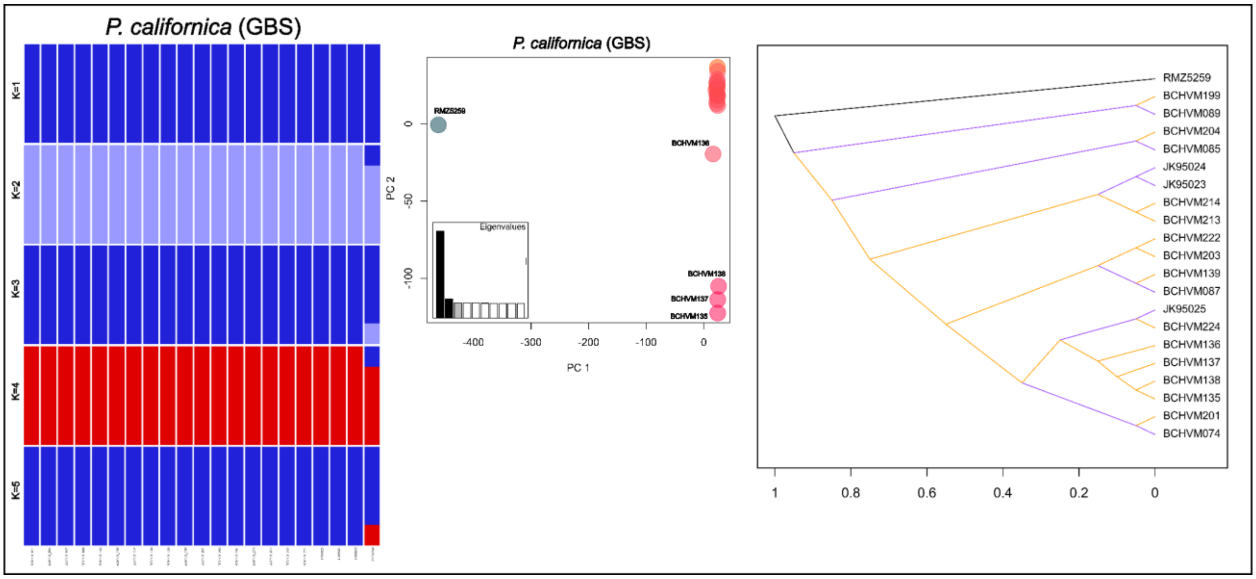


**Fig. S7.** Population clustering information for California Gnatcatcher (GBS). Structure plots (K=1-5), and SNP cloudogram, respectively. PCA points labeled as individuals, when applicable, remaining unlabeled samples cluster too closely to be readily distinguishable. See methods for additional details.

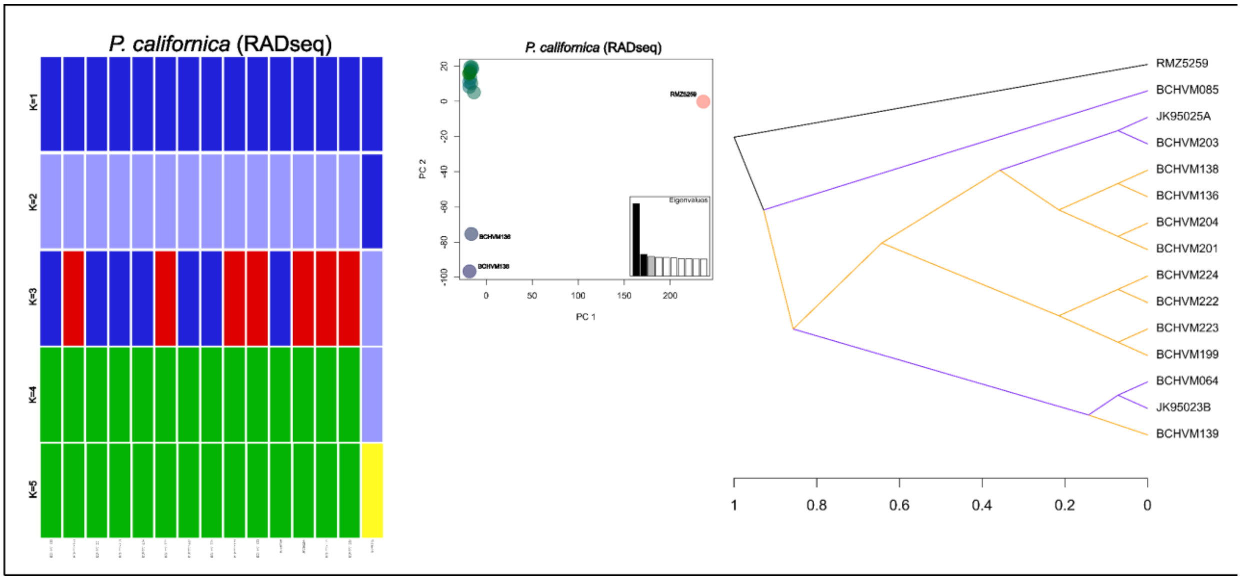


**Fig. S8.** Population clustering information for California Gnatcatcher (RADseq). Structure plots (K=1-5), PCA, and SNP cloudogram, respectively. PCA points labeled as individuals, when applicable, remaining unlabeled samples cluster too closely to be readily distinguishable. See methods for additional details.

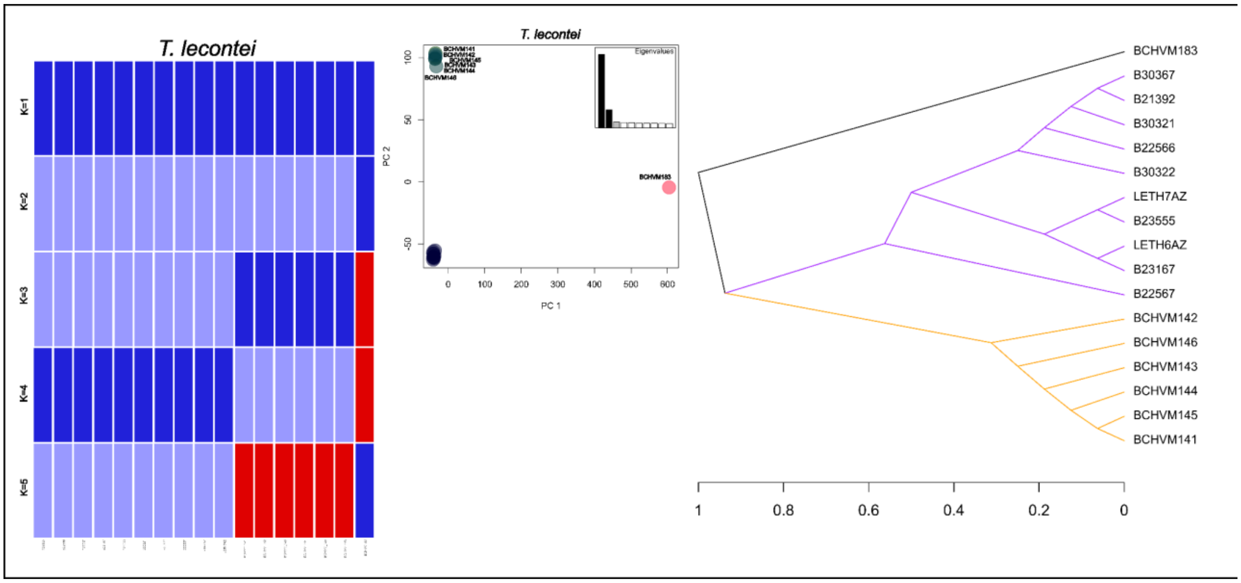


**Fig. S9.** Population clustering information for Le Conte's Thrasher (GBS). Structure plots (K=1-5), PCA, and SNP cloudogram, respectively. PCA points labeled as individuals, when applicable, remaining unlabeled samples cluster too closely to be readily distinguishable. See methods for additional details.
